# Stimulation augments spike sequence replay and memory consolidation during slow-wave sleep

**DOI:** 10.1101/670091

**Authors:** Yina Wei, Giri P Krishnan, Lisa Marshall, Thomas Martinetz, Maxim Bazhenov

## Abstract

Newly acquired memory traces are spontaneously reactivated during slow-wave sleep (SWS), leading to the consolidation of recent memories. Empirical studies found that sensory stimulation during SWS selectively enhances memory consolidation and the effect depends on the phase of stimulation. In this new study, we aimed to understand the mechanisms behind the role of sensory stimulation on memory consolidation using computational models implementing effects of neuromodulators to simulate transitions between awake and SWS sleep, and synaptic plasticity to allow the change of synaptic connections due to the training in awake or replay during sleep. We found that when closed-loop stimulation was applied during the Down states (90^0^-270^0^) of sleep slow oscillation, particularly right before transition from Down to Up state, it significantly affected the spatio-temporal pattern of the slow-waves and maximized memory replay. In contrast, when the stimulation was presented during the Up states (270^0^-360^0^ and 0^0^-90^0^), it did not have a significant impact on the slow-waves or memory performance after sleep. For multiple memories trained in awake, presenting stimulation cues associated with specific memory trace could selectively augment replay and enhance consolidation of that memory and interfere with consolidation of the others (particularly weak) memories. Our study proposes a synaptic level mechanism of how memory consolidation is affected by sensory stimulation during sleep.

**Significance statement:** Stimulation, such as training-associated cues or auditory stimulation, during sleep can augment consolidation of the newly encoded memories. In this study, we used a computational model of the thalamocortical system to describe the mechanisms behind the role of stimulation in memory consolidation during slow-wave sleep. Our study suggested that stimulation preferentially strengthens the memory traces when delivered at specific phase of slow oscillations just before Down to Up state transition when it makes the largest impact on the spatio-temporal pattern of sleep slow waves. In the presence of multiple memories, presenting sensory cues during sleep could selectively strengthen selected memories. Our study proposes a synaptic level mechanism of how memory consolidation is affected by sensory stimulation during sleep.

## Introduction

Memories depend on three general processes: encoding, consolidation and retrieval. After the new knowledge is encoded, it undergoes a phase of consolidation. Sleep facilitates this process by spontaneously reactivating the newly encoded memories (Rasch and Born, 2013; Born and Wilhelm, 2012; Diekelmann and Born, 2010b; Walker and Stickgold, 2004). A common hypothesis is that the consolidation of memories during sleep occurs through the reactivation of the neuron ensembles engaged during learning (Ji and Wilson, 2007b; Euston, Tatsuno and McNaughton, 2007; Peyrache *et al.*, 2009a; Barnes and Wilson, 2014; Ramanathan, Gulati and Ganguly, 2015).

Presently, sensory stimulation during non-rapid eye movement (NREM) sleep has been shown to modulate brain rhythms of sleep as well as affect memory performance across sleep. The sensory stimuli applied during sleep were either associated with the learning material in a train session before sleep - targeted memory reactivation (TMR) (Rudoy *et al.*, 2009; Rasch *et al.*, 2007; Antony *et al.*, 2012; Schonauer, Geisler and Gais, 2014; Cousins *et al.*, 2014; Cousins *et al.*, 2016), or were unassociated to learning content (Ngo *et al.*, 2013; Ngo *et al.*, 2015). In both cases sleep stimulation could improve memory performance compare to non-stimulation case (for review, see (Oudiette and Paller, 2013; Schouten *et al.*, 2017)). In TMR, auditory (Rudoy *et al.*, 2009) or olfactory (Rasch *et al.*, 2007) cues that were used during learning are presented again during NREM sleep leading to an increase in memory performance after sleep. While many TMR studies are employed for declarative memory tasks, TMR protocols may also enhance consolidation of hippocampus-independent procedural memories (Antony *et al.*, 2012; Schonauer, Geisler and Gais, 2014; Cousins *et al.*, 2014; Cousins *et al.*, 2016). Antony et al., (2012) trained participants for two different motor sequence tasks with sensory cues, and then presented the cue for one of the tasks during the nap. Following the nap, the cued task revealed higher performance than the uncued one, suggesting TMR can enhance procedural memories. The amount of improvement correlated with the duration of SWS and the number of spindles during SWS (Antony *et al.*, 2012). Similar improvements were found for the declarative component of a trained sequence (Cousins *et al.*, 2014). Memory performance across a sleep period of 8 hours without auditory cues was comparable to performance across a shorter sleep period of 3 hours with cued activation (Schonauer, Geisler and Gais, 2014). Auditory cues during SWS may not only enhance motor sequence performance, but also increase the functional brain activity and connectivity in consolidation-relevant networks (Cousins *et al.*, 2016). Interestingly, presentation of the cues during wakefulness was not effective for augmenting either declarative (Rasch *et al.*, 2007) or procedural memories (Schonauer, Geisler and Gais, 2014).

When the sensory stimuli were unassociated with the learning content, i.e. presenting pink noise during sleep, it was also found to enhance memory consolidation by enhancing the amplitude of slow waves during sleep (Ngo *et al.*, 2013; Ngo *et al.*, 2015). Stimulation was effective only when the sounds occurred in synchrony with the slow waves. In contrast, out-of-phase stimulation was ineffective (Ngo *et al.*, 2013). Surprisingly, in closed-loop auditory stimulation protocol, when more than two stimuli were presented sequentially at the adjacent cycles of the slow oscillation, it led to saturation and did not further enhance memory performance (Ngo *et al.*, 2015).

In this study, we used a biophysical model of the thalamocortical network to investigate the mechanisms behind the role of external stimulation - training-associated cues or sensory stimulation - on memory consolidation during SWS. The model incorporated populations of thalamic and cortical neurons and implemented effects of neuromodulators to allow transitions between awake and slow-wave sleep (SWS) states (Krishnan *et al.*, 2016), as well as STDP(spike-timing-dependent plasticity) type synaptic plasticity (Wei, Krishnan and Bazhenov, 2016; Wei *et al.*, 2018). Our study explains previous empirical data and provides insights into how synaptic reactivation within the thalamocortical network may be affected by the sensory stimulation during sleep.

## Materials and Methods

### Model description

#### Network geometry

The thalamocortical network model incorporated 40 thalamic relay (TC) and 40 reticular (RE) neurons in the thalamus, 200 pyramidal neurons (PY) and 40 inhibitory interneurons (IN) in the cortex (Wei, Krishnan and Bazhenov, 2016; Bazhenov *et al.*, 2002; Wei *et al.*, 2018) organized with local synaptic connectivity (Fig. 1). The PY and IN neurons received AMPA and NMDA synapses from PY neurons, and PY neurons also received GABA_A_ synapses from IN neurons. The radii of connections between cortical neurons were R_AMPA(PY-PY)_ = 5, R_NMDA(PY-PY)_ = 5, R_AMPA(PY-IN)_ = 1, R_NMDA(PY-IN)_ = 1 and R_GABAA(IN-PY)_ = 5. The TC neurons projected to RE neurons through AMPA synapses (R_AMPA(TC-RE)_ =8), and connections from RE to TC neurons included GABA_A_ and GABA_B_ synapses (R_GABAA(RE-TC)_ = 8, R_GABAB(RE-TC)_ = 8). The radii of connections between RE and RE were R_GABAA(RE-RE)_ = 5. Thalamocortical connections were wider and mediated by AMPA synapses from TC neurons (R_AMPA(TC-PY)_ =20, R_AMPA(TC-IN)_ = 4); corticothalamic connections were mediated by AMPA synapses from PY neurons (R_AMPA(PY-TC)_ =10, R_AMPA(PY-RE)_ =8). Flat connectivity profile was used for all synaptic connections. We previously tested different radii of connections and exponentially decaying profile and found qualitatively similar network dynamics, assuming that synaptic connections are scaled to maintain total synaptic input per neuron. All neurons were modeled based on the Hodgkin-Huxley kinetics. The units and description of parameters are summarized in Table 1.

**Figure 1.**
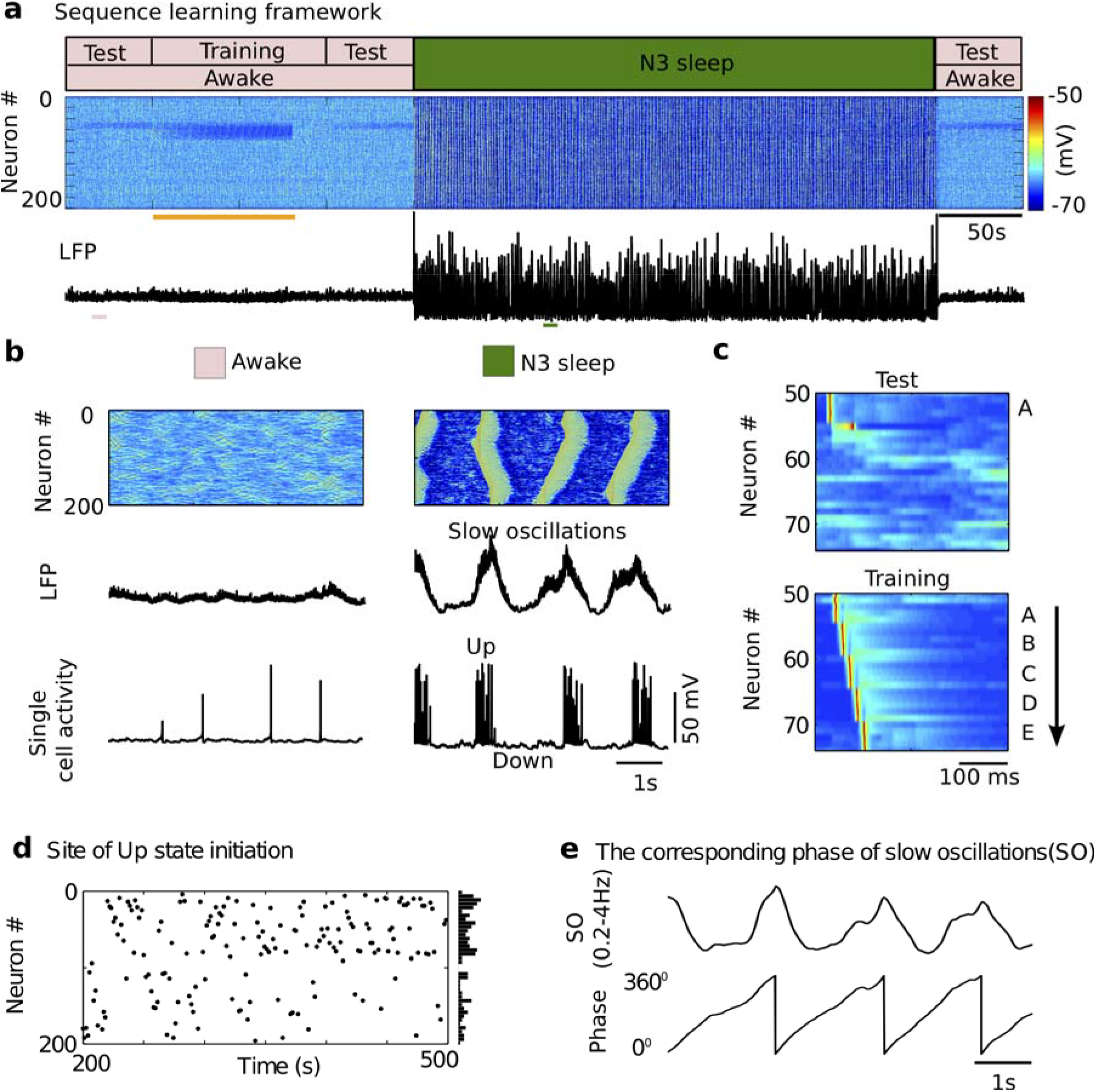
Network dynamics and sequence learning paradigm in the model. **a)** The cortical network activity during transitions from the awake state (pink block, *top*), to N3 sleep (dark green block) and back to awake. Raster plot (*middle*) shows membrane voltages of cortical pyramidal cells. Local field potential (LFP, *bottom*) from the cortical population of pyramidal cells. The sequence was learned during the training period (80s, orange bar). Performance was tested in three test sessions: before training, after training before sleep, and after sleep. **b)**. The expanded view of characteristic spatio-temporal patterns (*top*), LFP (*middle*) and single cell activity of neuron #50 (*bottom*) during awake (*left*), and N3 sleep (*right*) from where pink, dark green bars are shown under LFP traces in **a** (*bottom*). The slow oscillations (<1Hz) during N3 sleep consisted of a typical Up and Down state transitions. **c)** The characteristic examples of a training session and a test session. The training included stimulating sequentially (each group was stimulated for 10 ms and the delay between groups was 5 ms) at groups A, B, C, D, and E. The test included stimulating only at group A to recall the trained sequence within a 350ms response window. The sequence started at neuron #50. Each group included five neurons. **d)** Up state initiation sites over the entire sleep period are indicated by black dots. The vertical panel at the right represents probability of Up state initiation across neurons. **e)** The corresponding relationship between slow oscillations (SO) and phase.

**Table 1.**
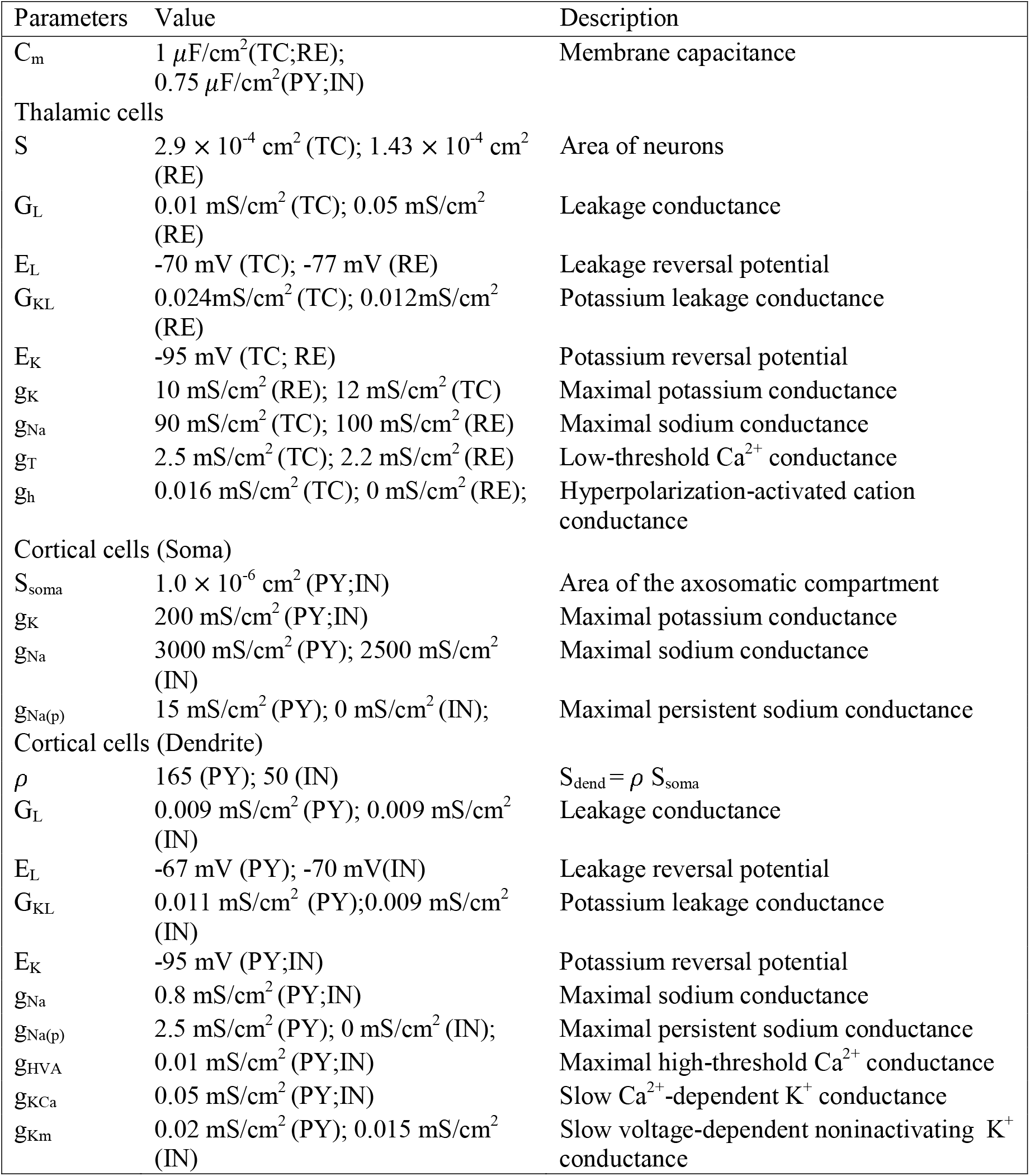
Main parameters. This table includes the units and description of the parameters used in the model.

#### Neuromodulators and sleep stages

The model implemented the change of neuromodulators, such as acetylcholine (ACh), histamine (HA), and GABA, in the intrinsic and synaptic currents to model transitions between sleep stages (Krishnan *et al.*, 2016). Specifically, the reduction of ACh was implemented as an increase of potassium leak conductance in TC, PY and IN neurons, a reduction of potassium leak conductance in RE cells (McCormick, 1992), and an increase in AMPA connection strength (Kimura, Fukuda and Tsumoto, 1999). The reduction of HA was implemented as a negative shift in the activation curve of a hyperpolarization-activated cation current (*I_h_*) (McCormick, 1992; McCormick and Williamson, 1991). The increase of GABA was implemented as an increase of the maximal conductance of the GABAergic synapses in IN and RE neurons (Krishnan *et al.*, 2016). These synaptic and intrinsic changes were tunes to model transitions between awake state and slow wave sleep (N3 stage) (Krishnan *et al.*, 2016).

#### Intrinsic currents: cortex

The cortical PY and IN neurons included dendritic and axo-somatic compartments, similar to the models used in our previous papers (Bazhenov *et al.*, 2002; Wei, Krishnan and Bazhenov, 2016; Timofeev *et al.*, 2000; Chen *et al.*, 2012; Krishnan *et al.*, 2016; Wei *et al.*, 2018), that is a reduction of the multi-compartmental neuron model as described in (Mainen and Sejnowski, 1996):

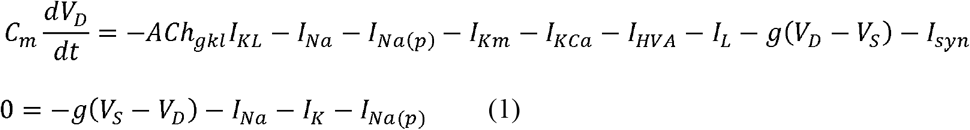

where *C_m_* is the membrane capacitance, *ACh_gkl_* represents the modulation on potassium leak current *I_KL_* based on the level of ACh during different sleep stages (ACh_gkl_=0.133 and 0.361 for awake and N3 sleep, respectively), *I_Na_* is a fast sodium current, *I_Na(p)_* is a persistent sodium current, *I_Km_* is a slow voltage-dependent non-inactivating potassium current, *I_KCa_* is a slow Ca^2+^-dependent K^+^ current, *I_HVA_* is a high-threshold Ca^2+^ current, *I_L_* is the Cl^−^ leak current, g is the conductance between axo-somatic and dendritic compartment. *V_D_* and *V_S_* are the membrane potentials of dendritic and axosomatic compartments, and I*_syn_* is the sum of synaptic currents to the neuron. This model was first proposed in (Mainen and Sejnowski, 1996) as a reduction of a multi-compartmental pyramidal cell model, based on the assumption that the current dynamics in the axosomatic compartment are fast enough to ensure that VS is always at equilibrium state, as defined by the second equation in Eq.(1). Indeed, this reduced model has relatively high Na^+^ and K^+^ conductance values (*g*_Na_ = 3000 mS/cm^2^, g_K_ = 200 mS/cm^2^ (Mainen and Sejnowski, 1996)) in the axosomatic compartment (representing axon hillock in the model). Therefore, the full version of the axosomatic membrane voltage equation *CdVs/dt* = −*g*(*V*_S_ − *V*_D_)– *I*_S_^int^ can be rewritten in a form. *ε* dVs/dt = F(Vs), where *ε* is a small parameter and F(Vs) represents axosomatic currents normalized to match the magnitude of the dendritic currents. Using singular perturbations analysis (Kuznetsov, 1995), we can find that the state variable *Vs* quickly reaches the manifold of slow motion defined by equation *F(Vs)=0*, that corresponds to Eq. (1) in our model. (See detailed discussion in (Chen *et al.*, 2012)). The persistent sodium current *I_Na(p)_* was included in the axosomatic and dendritic compartment of PY cells to increase bursting propensity. IN cells had the same intrinsic currents as those in PY except that *I_Na(p)_* was not included. All the voltage-dependent ionic currents *I_j_* have the similar form

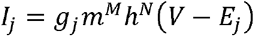

where g_j_ is the maximal conductance, m and h are gating variables, V is the voltage of the corresponding compartment and E_j_ is the reversal potential. The dynamic of gating variables are described as

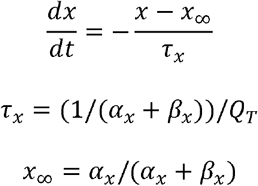

where x = m or h. Q_T_ is a temperature related term. Usually, Q_T_ = Q^((T-23)/10)^ =2.9529, with Q=2.3,T=36. The detailed description of individual currents was provided in our previous study (Wei, Krishnan and Bazhenov, 2016).

#### Intrinsic currents: thalamus

The thalamic TC and RE cells were modeled as a single compartment that included voltage- and calcium-dependent currents described by Hodgkin-Huxley kinetic (Bazhenov *et al.*, 2002):

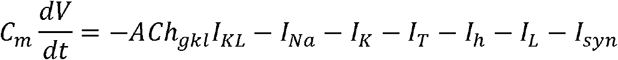

where *ACh_gkl_* in TC cells is 0.4, and 1.6 for awake, and N3 sleep. *ACh_gkl_* in RE cells is 0.9, and 0.45 for awake, and N3 sleep. *I_KL_* is a potassium leak current, *I_Na_* is a fast sodium current, *I_K_* is a fast potassium current, *I_T_* is a low threshold Ca^2+^ current, *I_h_* is a hyperpolarization-activated cation current, *I_L_* is a Cl^−^ leak current, and *I_syn_* is the sum of the synaptic currents to the neuron. The hyperpolarization-activated cation current *I_h_* was only included in TC neurons, not in RE neurons. The detailed description of individual currents was provided in our previous study (Wei, Krishnan and Bazhenov, 2016). The effect of HA on *I_h_* was implemented as a shift of *HA_gh_* in the activation curve (Krishnan *et al.*, 2016):

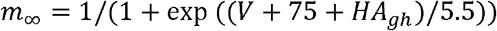

where *HA_gh_* is −24 mV, −1mV for awake, N3 sleep, respectively.

#### Synaptic currents

The equations for GABA_A_, AMPA, and NMDA synaptic currents were described by first-order activation schemes, and the GABA_B_ synaptic currents had a more complex scheme of activation that involved the activation of K^+^ channels by G proteins (Destexhe *et al.*, 1996). The equations for all synaptic currents used in this model were given in our previous studies (Wei, Krishnan and Bazhenov, 2016; Bazhenov *et al.*, 2002). In this paper, we added the level of ACh and GABA to modulate AMPA, and GABA_A_ synaptic currents as described by

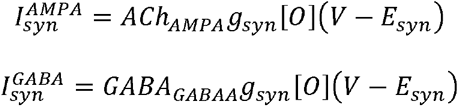

where g_syn_ is the maximal conductance, [O] is the fraction of open channels, and E_syn_ is the reversal potential (E_AMPA_=0 mV, E_NMDA_=0 mV, and E_GABAA_=-70 mV). ACh_AMPA_ is the variable that modulates AMPA synaptic currents for cortical PY-PY, TC-PY, and TC-IN connections by the level of ACh. ACh_AMPA_ from PY cells is 0.133, and 0.4332 for awake, and N3 sleep. ACh_AMPA_ from TC cells is 0.6, and 1.2 for awake, and N3 sleep. GABA_GABAA_ is the variable that modulates GABA synaptic currents for cortical IN-PY, RE-RE and RE-TC connections. GABAGABAA from IN cells is 0.22, and 0.44 for awake, and N3 sleep. GABA_GABAA_ from RE cells is 0.6, and 1.2 for awake, and N3 sleep, respectively.

The maximal conductance for each specific synapse was g_GABAA(RE-TC)_ = 0.06 μS, g_GABAB(RE-TC)_=0.0025 μS, g_GABAA(RE-RE)_ = 0.1μS, g_AMPA(TC-RE)_ = 0.06 μS, g_AMPA(TC-PY)_ = 0.14 μS, g_AMPA(TC-IN)_ = 0.12 μS, g_AMPA(PY-PY)_ = 0.24 μS, g_NMDA(PY-PY)_ = 0.01 μS, g_AMPA (PY-IN)_ = 0.12 μS, g_NMDA(PY-IN)_ = 0.01 μS., g_AMPA (PY-TC)_ = 0.06 μS, g_AMPA (PY-RE)_ = 0.1 μS and g_GABAA(IN-PY)_ = 0.24 μS.

In addition, spontaneous miniature EPSPs and IPSPs were implemented for PY-PY, PY-IN and IN-PY connections. The arrival times of spontaneous miniature EPSPs and IPSPs were modeled by Poisson processes (Stevens, 1993), with time-dependent mean rate μ = (2/(1+exp(-(t-t_0_)/F))-1)/250(Bazhenov *et al.*, 2002), where t0 is a time instant of the last presynaptic spike (Timofeev *et al.*, 2000). The mEPSP frequency (F) and amplitude (A) were F_PY-PY_ = 30, F_PY-IN_ = 30, F_IN-PY_ = 30, A_PY-PY_ =0.2 mV, A_PY-IN_=0.2 mV, and A_IN-PY_=0.2 mV.

#### Spike-timing dependent synaptic plasticity (STDP)

Facilitation or depression of the synaptic strength is believed to underlie learning in the brain. Here we used STDP model of synaptic plasticity to adjust the synaptic connections between cortical pyramidal neurons based on the relative timing of the pre- and postsynaptic spikes. The change of excitatory synaptic connections (g_AMPA_) and the amplitude of mEPSC (A_mEPSC_) were described as in our previous paper (Wei, Krishnan and Bazhenov, 2016):

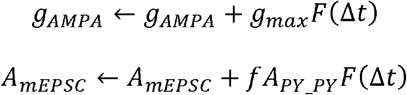

where g_max_ is the maximal synaptic conductance of g_AMPA_. f =0.01 is a factor representing the change of STDP on A_mEPSC_ is slower than on g_AMPA_. F is the STDP function that shows the change of synaptic connections as a function of the relative timing (Δt) of pre- and postsynaptic spikes (Song, Miller and Abbott, 2000),

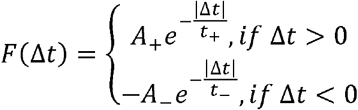

where parameters A_+_ and A_−_ determine the maximum amounts of synaptic modification. Here, we set A_+_ = A_−_ = 0.002, and *τ*_+_ = *τ*_−_ = 20 ms. We reduced the STDP amplitude A_+_ and A_−_ to 0.001 during slow-wave sleep to account for reduction of ACh (Sugisaki *et al.*, 2015). We assumed that the synaptic efficacy should stay within [0, 200%] range of the initial synaptic weights to prevent STDP from runaway synaptic dynamics. We would like to note that *in vivo* the rate of synaptic potentiation is slower than that in the model and typically saturates around 150% of cortical neurons over a full night (Chauvette, Seigneur and Timofeev, 2012). Because of that, although our simulation times (in absolute unites) are much shorter than a full night, the change of the synaptic weights in the trained region was sufficient to observe the performance improvement after sleep.

#### Training and Test

For most of the simulations, training pattern included 5 groups of neurons that were activated in sequential order in space and time, with 5 ms delay between subsequent groups activation. Each group was a set of 5 adjacent neurons drawn from a contiguous 25-cell subregion of the full 200-cell network. For example, if the sequence started at neuron #50, these 5 groups were: A(#50-54), B(#55-59), C(#60-64), D(#65-69),E(#70-74). Each group was stimulated by a step current that leads to a suprathrehold response with duration of 10 ms and a delay of 5 ms between groups. Thus during training, the neuronal activity in these groups reflected the order of the trained sequence, e.g., “ABCDE”. During test sessions, the model was only presented with the first input at group “A” to recall the trained sequence “ABCDE” within a 350ms response window. During both training and test sessions, each trial was repeated every 1s.

#### Stimulation

To model sensory cues, we modeled independently a second (sensory) cortical network with 200 pyramidal cells that one-to-one connected to the pyramidal cells of the primary network. The stimulation applies at the cue Q(#50-54) of the secondary cortical network which activates A(#50-54) of the primary network. During training session we stimulated sequentially Q→A→B→C→D→E. During sleep, in the open-loop stimulation protocol, the cue was triggered for 50ms at frequency 0.75Hz that was close to the internal frequency of slow oscillations in the model. In the closed-loop stimulation protocol, we first detected the onset of the Down state or Up state in ongoing slow oscillation (SO), then presented the cue with X ms delay after Down or Up state onset. We set X to be 0, 100, 200, 300, 400, or 500 ms. The Up or Down states were detected based on the analysis of the local field potential (LFP). The LFP was calculated by adding up membrane potentials of all the neurons, and it had a bimodal distribution (one peak corresponding to the Up states, and another peak corresponding to the Down states). The trough of the distribution was selected as a threshold to separate Up and Down state. The onset of Up or Down state was then defined as the moment when LFP value crossed the threshold. In the closed-loop “Best Phase” protocol, the stimulation was delivered at the optimal phase, which is ~500 ms after detecting the onset of a Down state, at each cycle of SO. In the closed-loop “2-Click” protocol, the stimulation was delivered at the optimal phase at two sequential cycles of SO with a pause of 2.5s after the second stimulation to match experimental protocol (Ngo *et al.*, 2015).

#### Computational methods

All model simulations were performed using a fourth-order Runge-Kutta integration method with a time step of 0.02 ms. Source C++ was compiled on a Linux server using the g++ compiler. Part of the simulation was run on the Neuroscience Gateway(Sivagnanam *et al.*, 2013). All data processing was done with custom-written programs in Matlab (MathWorks, Natick, MA).

### Data Analysis

#### Sequence learning analysis

To model sequence learning, the model was presented with multiple trials of sequential input to the groups of selected cortical neurons. The performance of sequence recall was measured by the percentage of success of sequence recall during test sessions when only the first group of a sequence was stimulated. First, we detected the network sequence using the following steps: 1) We detected all spikes for five groups of neurons (each group contains five neurons) within a 350ms response time window (starting from the time when test stimulus was applied); 2) We smoothed the firing rate of each group by convoluting the average instantaneous firing rate of five neurons with a Gaussian kernel (50ms window size); 3) The firing sequence of the groups was determined by ordering the peaks of their smoothed firing rates during 350ms window. Next, we applied a String Match (SM) method to measure the similarity between each detected sequence and an ideal sequence (e.g. S=“ABCDE”). The SM was calculated as 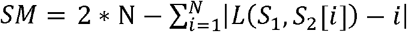, where *S*_1_ is the test sequence generated by the network, *S*_2_ is the subset of ideal sequence that only contains the same elements as *S*_1_, *N* is the sequence length of S_1_, *L*(*S*_1_,*S*_2_[*i*]) represents the location of element S_2_[*i*] in a sequence *S*_1_. SM was then normalized by dividing by M, where M is two times the length of S. For example, if the ideal sequence S was “ABCDE” and S_1_ was “ACDB”, then S_2_=“ABCD”, N =4. The location of element ‘A’ in S_1_ is L(S_1_, ‘A’)=1. ‘B’ in S_1_ is L(S_1_, ‘B’)=4, ‘C’ in S_1_ is L(S_1_, ‘C’)=2, ‘D’ in S_1_ is L(S_1_, ‘D’)=3. Therefore, SM = 2*4- (|1-1|+|4-2|+|2-3|+|3-4|)=4. After SM was normalized by M=10, it became 0.4, indicating the recalled sequence has 40% similarity to the ideal sequence. If the ideal sequence S was “ABCDE” and S_1_ was “ABCDE”, then S_2_=“ABCDE”, N =5 and SM=2*5-0=10, or 1.0 after normalization by 10. The performance was calculated as the percentage of recalled sequences with SM≥Th during the test session. In this paper, we selected a threshold of Th=0.8, indicating a recalled sequence with at least 80% similarity to the ideal sequence was counted as a successful recall. Baseline performance (before training) of the network was around 15% for Th=0.8 due to the random spiking. If higher threshold Th=1.0 was selected, the baseline performance became almost zero.

#### Analysis of synaptic weights

Synaptic weights between neurons in a direction of sequence activation were enhanced due to the sequence replay. The mean of the changes of synaptic weights associated with a given sequence was used to characterize memory strength.

### Statistical Analysis

When data were normally distributed based on statistical test, the numerical values are given as mean ± SEM, where SEM is standard error of the mean. Otherwise, we used median ± interquartile range (IQR) to report the data. For each experiment, 20 simulations with different random seeds were performed. Data were first tested for normal distribution by the Anderson-Darling test, and if data had a normal distribution, the parametric test was used; otherwise, the equivalent nonparametric test was applied. If only two groups of data were compared, the two-sample t-test (parametric) or the Mann–Whitney U test (nonparametric) was used. When data were paired, nonparametric Wilcoxon signed rank test was used. When more than two groups of data were compared, One-way ANOVA (parametric) or Kruskal-Wallis ANOVA test (nonparametric) with Bonferroni’s post hoc test was applied. To compare the means of two or more columns and two or more rows of the observations, two-way ANOVA was used.

## Results

### Open-loop periodic stimulation during sleep enhances memory performance

The thalamocortical network model was designed to model transitions between awake and slow-wave (N3) sleep (Fig. 1a) due to the changes in neuromodulators (Krishnan *et al.*, 2016). The characteristic network activity during the awake state (Fig. 1b, *left*) reveals random spiking and fluctuations in the local field potentials (LFP), while during N3 sleep the network displays slow (<1 Hz) oscillations (Fig.1b, *right*) characterized by repetitive transitions between Up and Down states in all cortical neurons (Steriade, Nunez and Amzica, 1993; Blake and Gerard, 1937; Steriade, Timofeev and Grenier, 2001). Similar to (Wei *et al.*, 2018), the awake state includes one training and three test sessions: before training, after training before sleep, and after sleep (Fig. 1a). During the training session, the model was presented with a sequence of stimuli applied to selected groups of cortical neurons (Fig. 1c, *bottom*). Each group contained five neurons - A(#50-54), B(#55-59), C(#60-64), D(#65-69), E(#70-74) – which were stimulated by 10 ms direct current steps that leads to a suprathrehold response. Five ms delay was included between stimulation of subsequent groups. Spike-timing dependent synaptic plasticity (STDP) during training changed synaptic connectivity. In result, the model learned a sequence, e.g., “ABCDE”. During test sessions, the model was only presented with the input at the first group “A” to test for pattern completion of the trained sequence “ABCDE” (Fig. 1c, *top*). During training or test sessions, each trial was repeated every 1sec. Initiation of Up states was random in the naive network and was biased to the trained network site after sequence learning (Fig. 1d). The instantaneous phase of the ongoing slow oscillations was derived from a Hilbert transformation of the simulated local field potentials (LFP): the peak of the Up state is defined as 0 degrees after Hilbert transformation, and the peak of the Down state is 180 degrees (Fig. 1e).

To model sensory cues, we modeled independently a second (“sensory”) cortical network making one-to-one connections to the primary network. Thus, during the training session we stimulated sequentially Q→A→B→C→D→E, where group of neurons (Q) belonged to the “sensory” network. During sleep, in an open-loop stimulation protocol, the cue was triggered for 50 ms at a pre-defined frequency 0.75Hz that was close to the internal frequency of slow oscillations in the intact model (Fig. 2a). The cue (Fig. 2b, *top*, red arrow) could trigger ABCDE sequence replay (Fig. 2d) and, if it occurred during later part of the Down state, it could also initiate an Up state of the slow oscillation (Fig. 2b, *bottom*).

**Figure 2.**
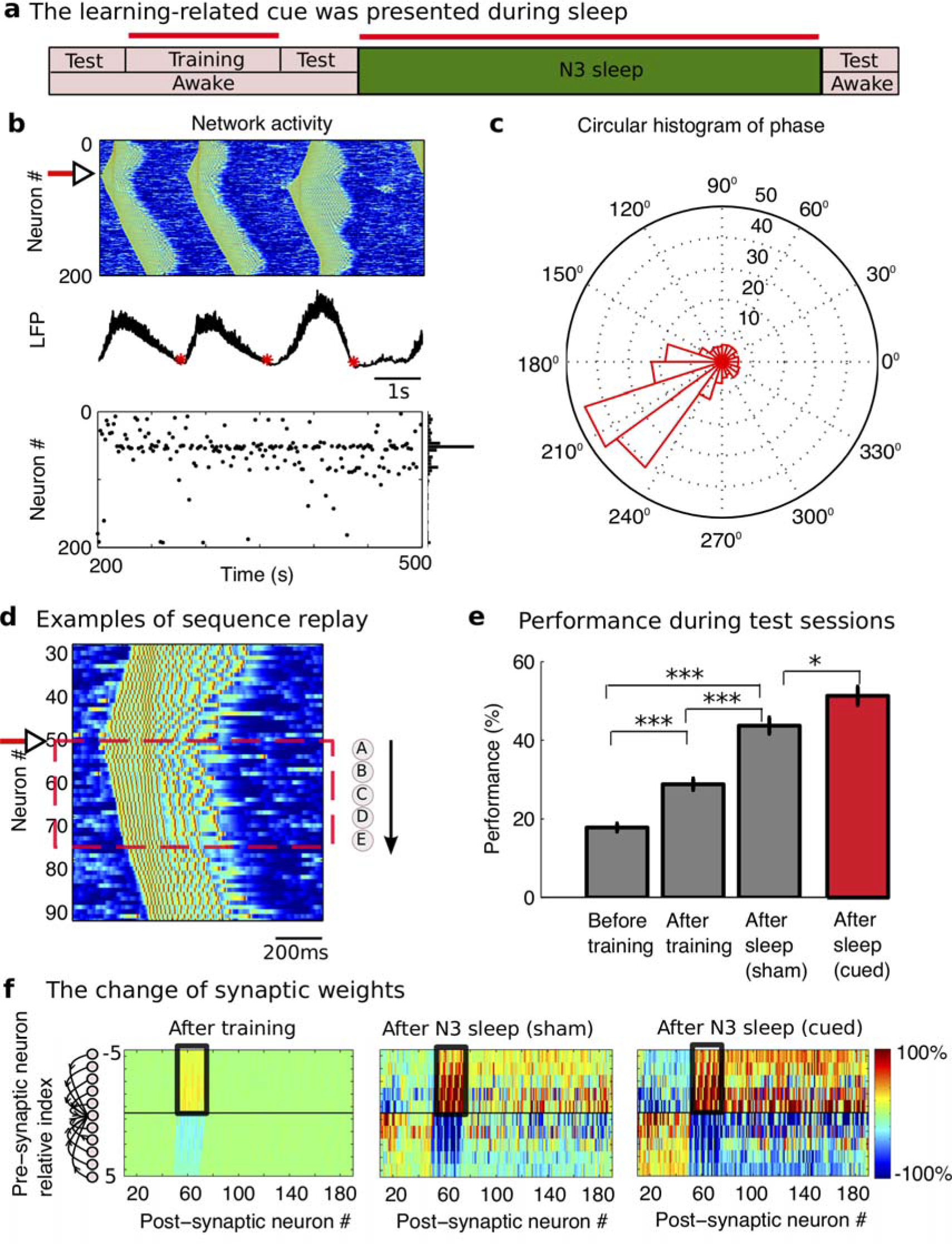
Periodic cue presentation during sleep. **a)** The cue was associated with the sequence during training session. During sleep, this learning-related cue was periodically presented for 50ms at every 1.333s (around 0.75Hz) during the whole sleep period. Red lines represent when the cue was presented. **b)** *Top*, Characteristic example of network activity during slow oscillations. Red arrow indicates cue input to group A. *Middle*, Characteristic example of LFP. The red dots indicate the times of cue presentation. *Bottom*, Up state initiation sites over the entire sleep period are indicated by black dots. The vertical panel at the right represents probability of Up state initiation across neurons. **c)** Circular histogram of slow oscillation phases at which the cue was applied. **(d)** Characteristic examples of sequence (“ABCDE”) replay during slow oscillations when the learning-related cue was presented. **e)** The bar plot of the performance that was defined by the probability of the recalled sequence with 80% similarity to the ideal sequence “ABCDE” (SM>=0.8) as measured during each recalled test session. Error bars indicate standard error of the mean (SEM). * p<0.05, ** p<0.01, *** p<0.001. **f)** The change of synaptic weights relative to the initial values after training (*left*): after N3 sleep without cue presentation (*middle*) and after N3 sleep with periodic cue presentation (*right*). The x-axis gives the index of the postsynaptic neuron; the y-axis represents the left (−) and right (+) presynaptic neurons relative to the postsynaptic neurons at x-axis. The synaptic weights between neurons in direction of sequence activation (black box) were enhanced due to training during awake (left) and sequence replay during sleep (right).

The phase of the slow oscillation at the time of a cue presentation (Fig. 2b, *middle*, red dots) was extracted to construct the circular histogram of the stimulation phase. The distribution had a well-formed peak (Fig. 2c) indicating that spontaneous slow oscillation was entrained or phase locked to the periodic cue stimulus. Analysis of the sequence recall performance among all three test sessions (Fig. 2e, grey bar) revealed the following (one-way ANOVA; F_2,57_=68.49, p=6.95*10^-16^). First, repetitive training for 80 sec during awake improved performance compared with the baseline performance before training (28.8%±1.454% vs. 17.8%± 1.025%, p=2.0677*10^-5^, oneway ANOVA Bonferroni corrections). Second, performance after sleep was enhanced significantly compared with that before sleep without cue stimulation (43.7%±2.059% vs. 28.8%±1.454%, p=2.926*10^-8^, one-way ANOVA Bonferroni corrections). Finally, when the cue was presented during sleep, performance after sleep (Fig. 2e, red bar) was significantly higher compared with the un-cued experiments (51.3%±2.314% vs. 43.7%±2.059%, t(38)= 2.454, p =0.0188, two-sample t-test).

To identify the change of the network connectivity underlying performance increase, we next analyzed the dynamics of synaptic weights between the cortical neurons. During the training phase, the ordered firing of neurons led to the potentiation of synapses in the direction of the trained sequence (Fig. 2f, *left*, black box), while the synapses corresponding to the opposite direction were depressed (Fig. 2f, *left*). These changes of the connectivity matrix among the trained neurons (Fig. 2f, *left*, black box) were augmented after the subsequent N3 sleep in both uncued (Fig. 2f, *middle*, black box) and cued experiments (Fig. 2f, *right*, black box), similar to our previous analysis of the replay without cues (Wei *et al.*, 2018). However, the changes were more significant in the experiments with sensory cue stimulation (Fig. 2f, *right*) which explains the higher performance of memory recall after the cued sleep compared to the uncued sleep.

The number of Up states of slow oscillation was significantly increased during cued sleep compared with uncued experiments (189.1±0.85 vs. 166.75±0.615, t(38)=-21.32, p =9.64*10^-23^, two-sample t-test, Fig3a red vs. black). This is consistent with the data of the recent experiments where the greater amount of slow oscillations was found to be elicited with the targeted sound compared to the experiments with a control new sound (Oyarzun *et al.*, 2017). To examine whether the enhanced performance in our simulations was merely due to the higher Up state count, we reduced the sleep duration from 300s to 260s for the cued sleep to obtain comparable Up state count to the uncued sleep. We observed that the performance after the 260 sec of cued sleep was still significantly higher than after the 300 sec of uncued sleep (50.6%±2.559% vs. 43.7%±2.059%, t(38)= −2.1, p =0.0423, two-sample t-test, Fig. 3a, pink vs. black). This suggests that the performance improvement due to cue presentation can be summed up in two factors: 1) facilitation of the trained sequence replay within each Up state; 2) increase in the number of Up state events.

**Figure 3.**
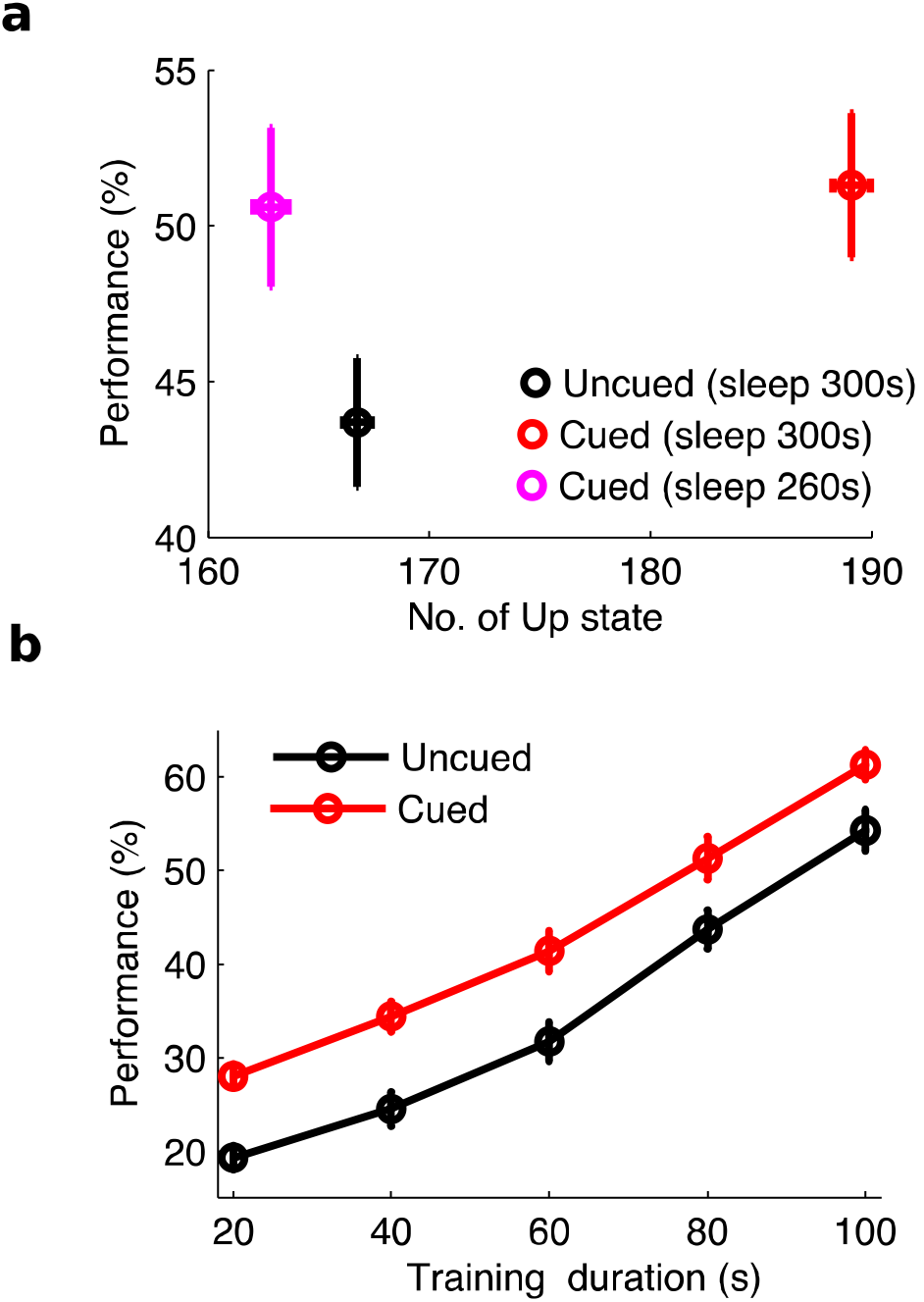
The effect of periodic cue presentation during sleep. **a)** The number of Up state during sleep vs. the performance after sleep. **b)** The performance after sleep for the different training duration (memory strength) with (red) or without cue during sleep (black). Error bars indicate SEM.

To characterize how relative strength of a memory trace before sleep influences the sequence recall and memory performance after the sleep, we varied duration of initial training (Fig. 3b). As the training duration (memory strength before sleep) increased, the performance after sleep increased as well (Fig 3b, compare among different memory strength, F(4,190)=106.33, p=2.2*10^-47^, two-way ANOVA). In all cases, performance of the sequence recall after sleep was significantly higher when the cue was presented during sleep (Fig. 3b, compare black and red, F(1,190)=51.60, p = 1.5*10^-11^, two-way ANOVA). Thus, the cues presented during sleep can benefit memory consolidation for different levels of the pre-sleep memory strength.

### Closed-loop stimulation protocol revealed phase-dependent response during SWS

Experimental studies suggest that stimuli presented at certain phases of the sleep slow oscillation have a stronger effect on memory consolidation compared to the other stimulation phases (Batterink, Creery and Paller, 2016). To investigate the relationship between the phase angle of the slow oscillation at the cue presentation and the change in performance, we used a closed-loop stimulation protocol to vary the timing of stimulation in respect to the phase of slow oscillation. In the closed-loop conditions, we first detected the onset of the Down state or Up state in ongoing slow oscillation, then presented the cue with X ms delay after Down or Up state onset. The onset of Up or Down state was defined as the moment when LFP crossed the threshold that was defined as the trough that separates two peaks in the histogram of the LFP distribution (see Methods). We set X to be 0, 100, 200, 300, 400, or 500 ms. In the example shown in Fig. 4a, the cue was presented with 500ms delay after the network transition from Up to Down state (Fig. 4a, *left middle*, red dot). After the experiment was completed, we plotted the circular histogram of stimulation phase for all stimulation events (Fig. 4a, right). In this particular example, the peak of the circular histogram was around 210^0^ (Fig. 4a, *right*), which corresponds to the very end of the Down state of the slow oscillation. We propose that since this was a time moment when the network was “almost ready” to start a new Up state on its own, stimulation reliably triggered a new Up state and, because of the stimulus location, these Up states were largely initiated near the beginning of the trained sequence (group “A”) thus promoting sequence replay (Fig. 4a, *left bottom*, see peak of histogram). In another characteristic example in the Fig. 4b, the cue was presented 200ms after the onset of Up state (Fig. 4b, *left middle*, red star). At that phase of stimulation, the entire network was already in the Up state (phase around 330^0^ (Fig. 4b, *right*)), and therefore stimulation had minimal impact on the network spatio-temporal dynamics (Fig. 4b, *left bottom*).

**Figure 4.**
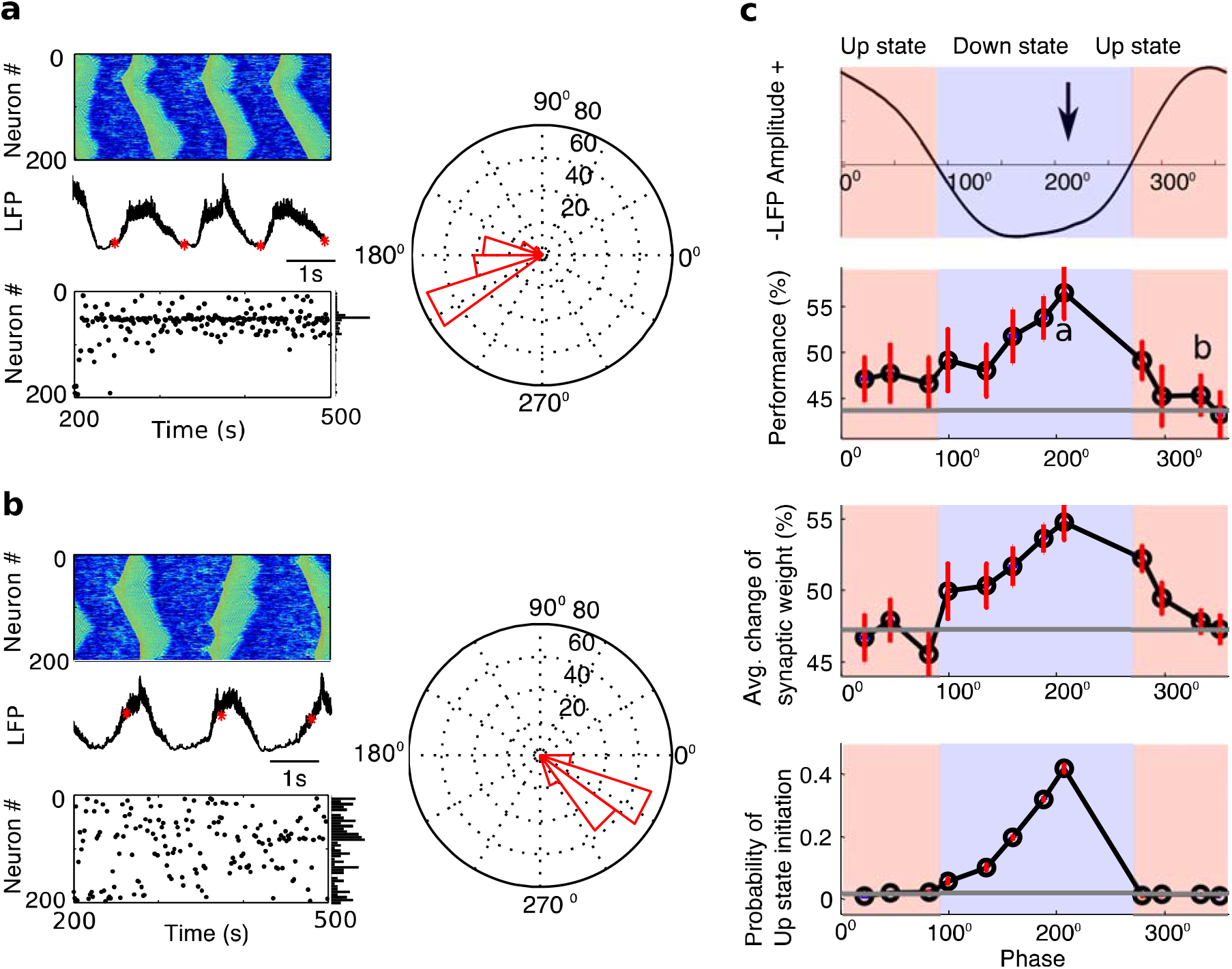
A closed-loop stimulation during slow oscillations. **a, b)** Examples of cue presentation during Down and Up state. The cue was presented in panel **a** and **b** correspond to 500ms after Down state started and 200ms after Up state started. In panel **a** and **b**, *Top left*, Characteristic example of network activity during slow oscillations. *Middle left*, Characteristic example of LFP. The red stars indicate the times of stimulation. *Bottom left*, Up state initiation sites over the entire sleep period are indicated by black dots. The vertical panel at the right represents probability of Up state initiation across neurons. *Right*, Circular histogram of phases at which the cue was applied. **c)** *Top*, the correspondence between phase angle and states of slow oscillation (Up and Down state). The vertical arrow indicates the optimal phase for cue presentation. *Middle Top*, the performance after sleep as a function of phase. The letter “a” and “b” correspond to the peak phase of panel **a** and **b**. *Middle bottom*, the change of synaptic weights associated with trained sequence as a function of phase. *Bottom*, the probability of Up state initiation at group A (neuron #50-54) as a function of phase. Error bars indicate SEM. Grey line represents the value from uncued sleep. The blue and red regions correspond to Down and Up states of slow oscillation respectively.

Fig 4 shows summary of results for all stimulation phases. When the cue was presented during the Down state (90^0^-270^0^) of slow oscillations, the performance (Fig. 4c, *middle top*), the synaptic weight change associated with the trained sequence replay (Fig. 4c, *middle bottom*), and the probability of Up state initiation at group “A” of neurons (Fig. 4c, *bottom*), were all higher than those when the cue was presented during the Up state (270^0^-360^0^ and 0^0^-90^0^), and also higher than those in the uncued model (Fig.4c, grey lines). The optimal phase was around 210^0^, which was right before transition from Down to Up state, and it maximized synaptic changes and performance improvement (Fig. 4c). It is worth noting that because of the relatively small size of the model, open-loop periodic stimulation at the frequency close to the natural frequency of slow waves entrained the entire network oscillation by selecting the same optimal phase of stimulation as we report here using closed-loop protocol (compare Fig. 2c and 4a).

Analysis of the Up state initiation sites (see examples in Fig. 4a,b, histograms) revealed that in the model with optimal stimulation phase (Fig. 4a) stimulation leads to the majority of Up states being initiated at the location corresponding to the beginning of the trained sequence, which promotes sequence replay. In contrast, for suboptimal stimulation time (Fig. 4b), distribution of the Up state initiation sites remains random. Our modeling findings are in agreement with recent experimental data that revealed that the learning-related cues would preferentially strengthen associated memories when they are delivered at the Down phase of slow oscillation (Batterink, Creery and Paller, 2016). Our study proposes a possible mechanism for such phase dependence – reorganization of the structure of the slow waves to promote specific sequence replay - and provides insight into how the memory consolidation may be affected by sensory cues applied during sleep slow oscillation.

### Closed-loop stimulation enhances SO power and peak frequency

To compare model predictions with experimental data (Ngo *et al.*, 2015; Ngo *et al.*, 2013), we next tested performance of the closed-loop “2-Click” protocol. Above, we found that the optimal phase for stimulation was when the cues were delivered ~500ms after detecting the onset of a Down state. In the “Best Phase” protocol stimulation was delivered at the optimal phase *at each cycle* of SO. The “2-Click” protocol consists of 2 subsequent clicks delivered at the optimal phase with a pause of 2.5s after the second click (Fig. 5a) to match experimental protocol (Ngo *et al.*, 2015). In the Sham (control) condition, no cue was presented during sleep. Compared with the Sham condition, the cue presentation at “Best Phase” significantly increased SO power (Fig. 5b, p=3.18*10^-13^, pairwise comparison), shifted the peak frequency in the SO range (0.2-1 Hz) towards higher frequencies (Fig. 5c, p=1.3*10^-11^, pairwise comparison), and increased the performance of sequence recall after sleep (Fig. 5d, p=0.001, pairwise comparison). These results are all consistent with experimental data (Fig.5 e,f,g, modified from Ngo et al. 2015). Surprisingly, overall effects of the “Best Phase” and “2-Click” stimulation protocols were very similar. Although there were slight differences in the slow oscillation power and peak frequency (Fig. 5b, p=0.0324; Fig. 5c, p=0.0104; pairwise comparison), there was no significant difference in performance between “Best Phase” and “2-Click” stimulation protocols after sleep (Fig. 5d, p=0.0773, pairwise comparison). Importantly, this result is consistent with experimental data (Fig. 5 e,f,g, modified from Ngo et al. 2015).

**Figure 5.**
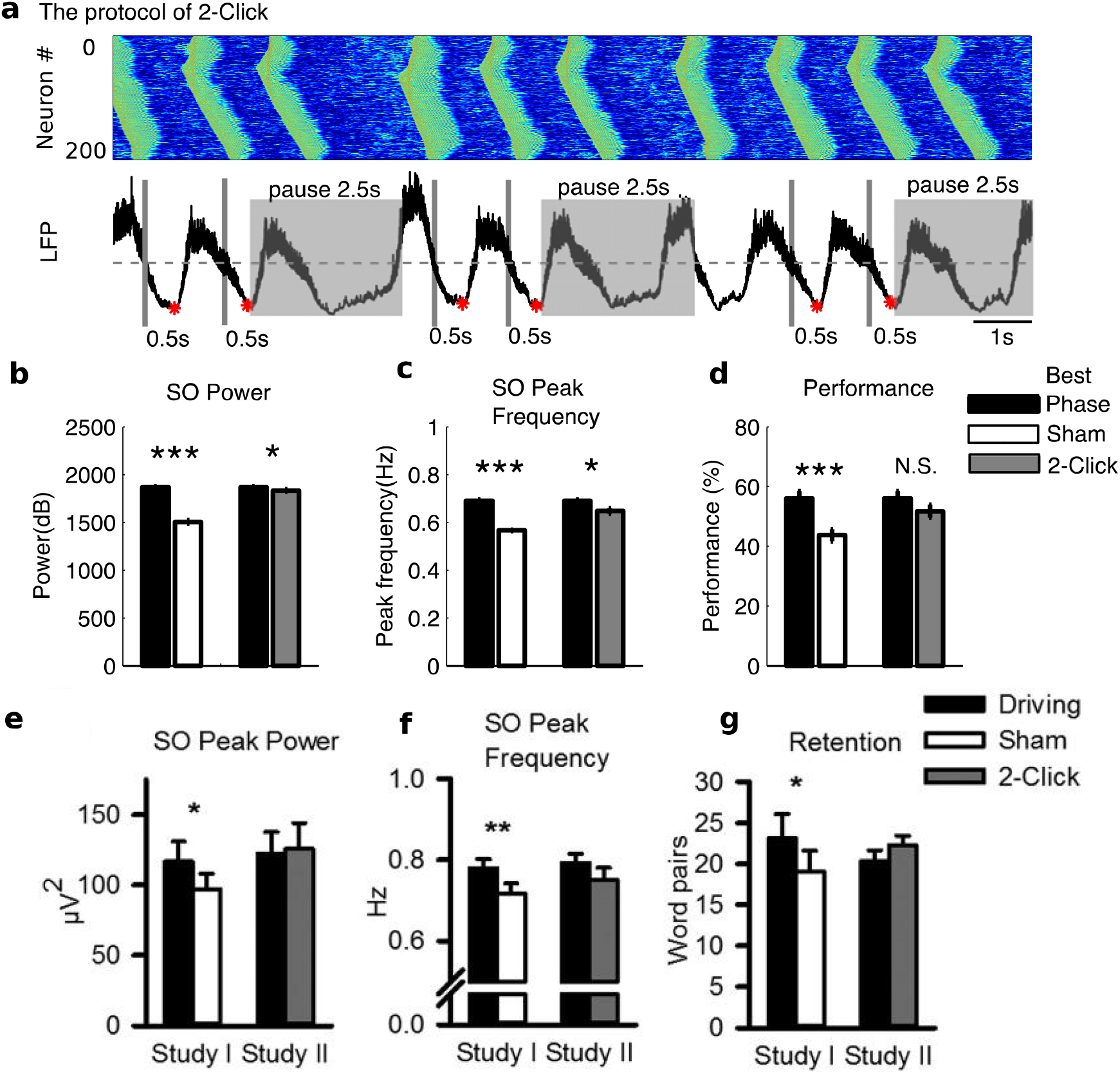
Closed-loop stimulation enhances SO power, peak frequency, and performance. **a)** The protocol of two clicks. The cue was triggered at 0.5s after detecting the onset of Down state. The detection routine was paused for 2.5s after the second cue presentation. Red star indicates the time when the cue was presented. **b,c,d)** The SO power, SO peak frequency and performance after sleep was compared between “Best Phase” and Sham, as well as between “Best Phase” and “2-Click”. The protocol of “Best Phase” was the same as “2-Click”, but without pausing the detection routine. For Sham condition, time points of stimulation were marked but no actual cue was presented. **e,f,g)** The SO peak power, SO peak frequency and retention from experimental data (modified from Ngo et al, 2015).

We previously reported that the spatio-temporal pattern of slow waves has a “history effect” – after a slow wave is initiated at one network site, there is a higher probability for initiation to occur again at the same site for 3-4 subsequent cycles (Wei, Krishnan and Bazhenov, 2016). We believe this phenomenon can explain the results of our new stimulation experiments. When only 1 or 2 cycles of slow oscillation were skipped between pairs of stimuli, the network still had higher than chance probability for Up state initiation at the previous initiation site (that was the first group of neurons in the trained sequence). Only when few cycles were skipped in a row, Up states began initiating randomly and this led to performance degradation.

### Closed-loop stimulation protocol with the entire network activation

From perspective of the cortical neurons, it remains unclear how focal or distributed the effect of the sensory stimulation is during sleep. In the next set of model stimulations, instead of the input targeting only a single group “A” of five neurons, stimulation targeted a much larger neuronal population (Fig. 6a,*left top*). The phase of closed-loop stimulation was selected near the end of a Down state, before the onset of an Up state (Fig. 6a, *right*), based on the results we obtained in the previous modeling experiments. We found that the amplitude of SO (Fig. 6a, *left middle*) was significantly increased in such stimulation conditions due to the very synchronized network transitions from Down to Up states (Fig. 6a, left). Up state initiation front was also broader (more neurons transited to Up states within relatively small time window), but still was centered near the trained population of neurons (Fig. 6a, *left bottom*). In another experiment (Fig. 6b), the broad stimulation cue was presented 200ms after the onset of Up state (Fig. 6b, *left middle*, red star). At that phase of stimulation, the large fraction of the cortical neuron population was in the Up state already (phase around 330^0^ (Fig. 6b, *right*)), and therefore stimulation had minimal impact on the location of the Up state initiation site and slow-wave spatio-temporal pattern (Fig. 6b, *left bottom*). Thus, when the broad cue was presented during Down state (90^0^-270^0^) of slow oscillations, the performance (Fig. 6c, *middle top*), the synaptic weight change associated with the sequence (Fig. 6c, *middle bottom*), and the probability of Up state initiation at group “A” of neurons (Fig. 6c, *bottom*), were all higher than those when the cue was presented during the Up state (270^0^-360^0^ and 0^0^-90^0^), and also higher than those in the uncued model (Fig. 6c, grey lines). These findings are similar to what we observed during the local cue presentation.

**Figure 6.**
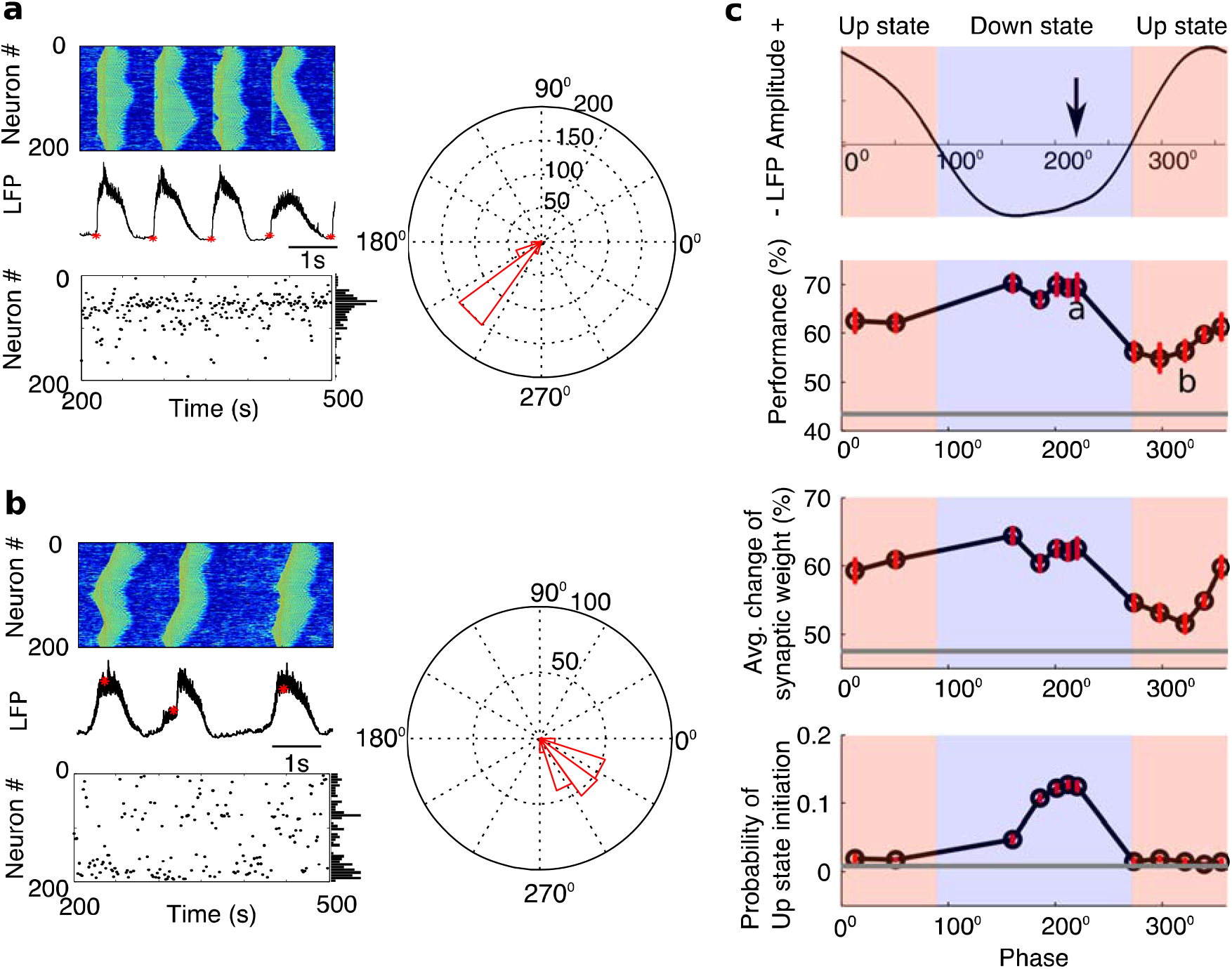
A closed-loop stimulation protocol with a broad cue input during slow oscillations. **a, b)** Examples of cue presentation during Down and Up state. The cue was presented in panel **a** and **b** correspond to 500ms after Down state started and 200ms after Up state started. In panel **a** and **b**, Top left, Characteristic example of network activity during slow oscillations. Middle left, Characteristic example of LFP. The red stars indicate the times of stimulation. Bottom left, Up state initiation sites over the entire sleep period are indicated by black dots. The vertical panel at the right represents probability of Up state initiation across neurons. Right, Circular histogram of phases at which the cue was applied. **c)** Top, the correspondence between phase angle and states of slow oscillation (Up and Down state). The vertical arrow indicates the optimal phase for cue presentation. Middle Top, the performance after sleep as a function of phase. The letter “a” and “b” correspond to the peak phase of panel **a** and **b**. Middle bottom, the change of synaptic weights associated with trained sequence as a function of phase. Bottom, the probability of Up state initiation at group A (neuron #50-54) as a function of phase. Error bars indicate SEM. Grey line represents the value from uncued sleep. The blue and red regions correspond to Down and Up states of slow oscillation respectively.

To quantify how external stimulation affects the spatiotemporal pattern of slow oscillation, we measured the phase locking index (PLI) of focal LFPs during individual Up states (Fig. 7a, each line represents the population activity from 25 neurons, we excluded the boundary neurons from analysis). The following three conditions were compared (Fig. 7b): Sham (PLI=0.4826±0.0306), cueing a local group of neurons (“local cue”) at the optimal stimulation phase (PLI=0.6387±0297), and cueing a broad group of neurons (“broad cue”) at the optimal stimulation phase (PLI=0.5793±0.0166). We found that PLI for the Sham group was significantly lower than for the model using local stimulation at site “A” only (p = 1.72*10^-25^, one-way ANOVA Bonferroni corrections) as well as for the model using broad stimulation (p=4.06*10^-16^, one-way ANOVA Bonferroni corrections).

**Figure 7.**
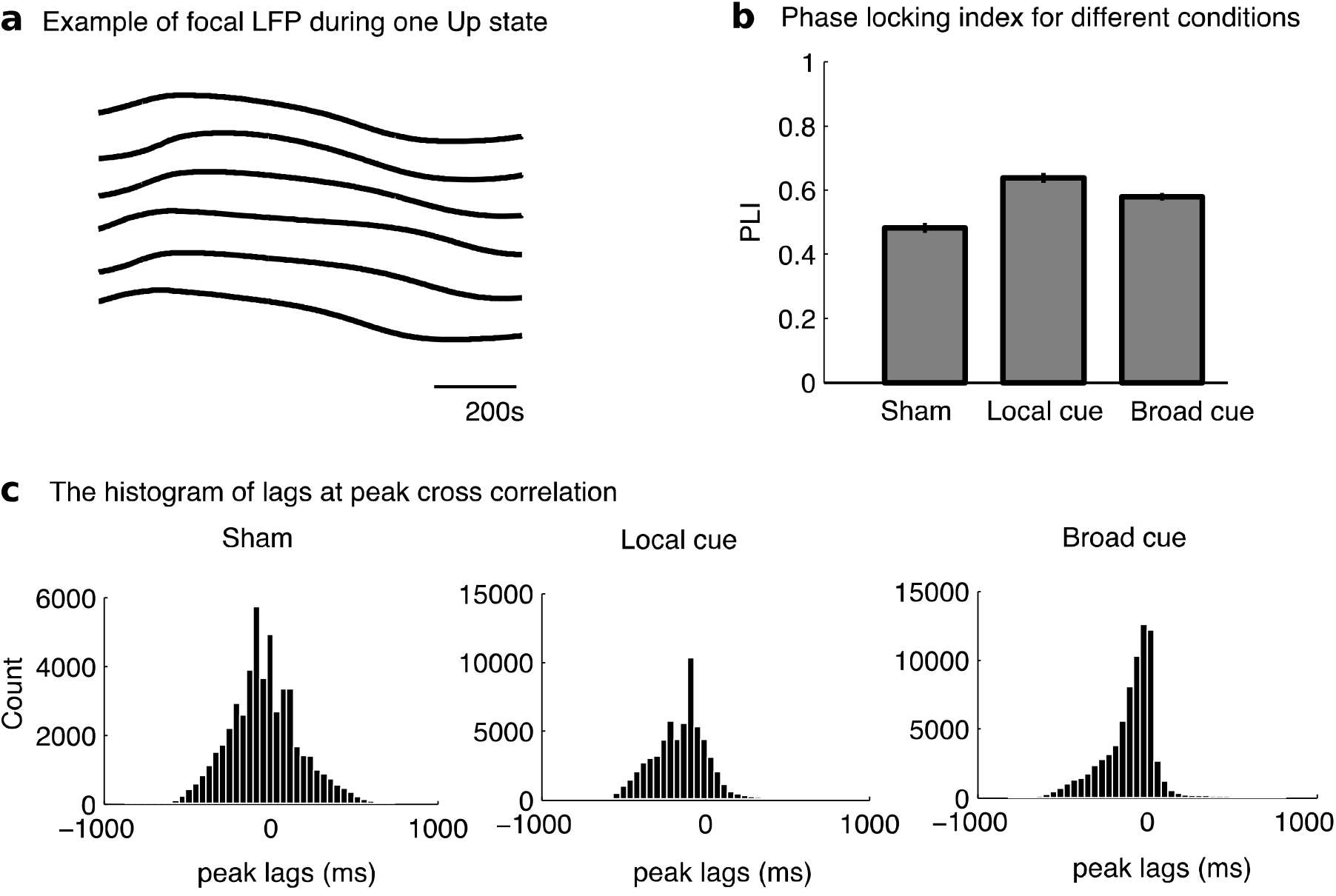
The spatial pattern of slow oscillations in three different conditions during sleep: Sham, local cue and broad cue. **a)** Example of population activity (every 25 neurons) during one Up state. The boundary neurons are not included. **b)** Phase locking index for different conditions: Sham, local cue and broad cue. Both local and broad cue was presented at the optimal phase. The local cue targeted only at group A, while the broad cue targeted at all the neurons excluding the boundary neurons. **c)** The histogram of lags at peak cross correlation at three conditions.

We next calculated cross correlations between all possible LFP pairs and plotted the distribution of the time lags to the peak (Fig. 7c). In Sham condition, the distribution was relatively broad with multiple local peaks (Fig. 7c, *left*) suggesting that many different network sites could lead transition from Down to Up state at different SO cycles or even any one cycle of SO. When the local stimulation was applied at the optimal phase, the lags peaked around −100ms (Fig. 7c, *middle*), suggesting the pattern of propagation of the slow waves from one preferential location, which can be explained by that the locally stimulated region was the main initiation site. When the stimulation was applied across a broad region at the optimal phase, the lags peaked around 0 ms (Fig. 7c, *right*), indicating that the broad stimulation increased network synchronization with zero lag.

### Selective stimulation targets specific memories during sleep replay

To test how the targeted stimulation during sleep affects only one memory when several memories are trained, we trained two independent sequences at two different network locations (Fig. 8a). We used long enough initial training time for both sequences to ensure that they replay and consolidate during slow-wave sleep regardless of the interference effects (Wei *et al.*, 2018). In the model, two sequences were trained by sequentially presenting stimuli at group A_1_(#50-54), B_1_(#55-59), C_1_(#60-64), D_1_(#65-69), E_1_(#70-74) for Seq1, and at group E_2_(#146-150), D_2_(#141-145), C_2_(#136-140), B_2_(#131-135), A_2_(#126-130) for Seq2, respectively (Fig. 8b, right). Each sequence was paired with a different sensory cue. During test sessions, as before, the recall performance for each sequence was measured based on the network response by stimulating only the first group of neurons in each sequence: group A_1_, or group E_2_ (Fig. 8b, *left*). When two sequences were trained for the same duration 80s, and no cues were presented during sleep, the recall performance was similar for both tasks (Fig. 8c and 7d, *left*, Uncued). Specifically, the performance for recalling Seq1 before training, after training before sleep, and after sleep were 17.9%±1.37%, 27.4%±1.43%, and 44.7% ± 2.02%, respectively (Fig. 8c, *top*), while the performance for recalling Seq2 before training, after training before sleep, and after sleep were 17.5%±0.96%, 32.6%±1.72%, and 49.8%±2.46%, respectively (Fig. 8c, *bottom*). For Seq1, the performance was significantly increased after training compared with before training (p=3.78*10^-4^, one-way ANOVA, Bonferroni corrections), and further significantly improved after sleep compared with before sleep after training (p=1.44*10^-9^, one-way ANOVA, Bonferroni corrections). For Seq2, the performance was significantly increased after training compared with before training (p=6.94*10^-7^, one-way ANOVA, Bonferroni corrections), and further significantly improved after sleep compared with before sleep after training (p=3.12*10^-8^, one-way ANOVA, Bonferroni corrections).

**Figure 8.**
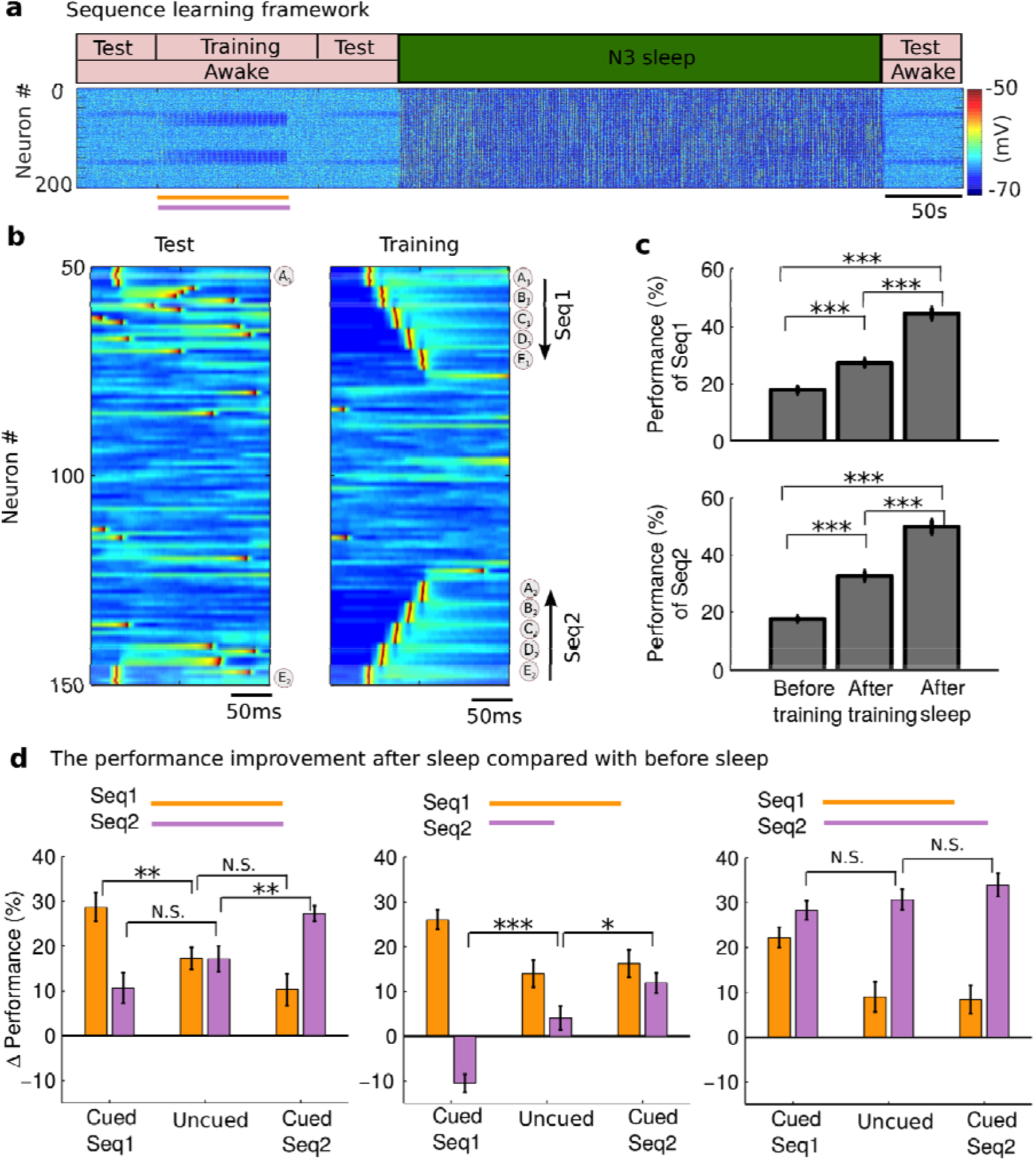
The cue can selectively enhance the specific memory. **a)** The model simulated transitions from awake to N3 sleep, and to awake again. Two sequences were trained during awake. Orange and purple bars represent the duration of training for Seq1 and Seq2, respectively. **b)** Characteristic example of test and training of Seq1 (“A_1_B_1_C_1_D_1_E_1_”) and Seq2 (“E_2_D_2_C_2_B_2_A_2_”). The test was stimulating only at group A_1_ for Seq1 and group E_2_ for Seq2. The Seq1 and Seq2 started at neuron #50 and #150, respectively. **c)** The bar plots of performance for Seq1 and Seq2 during test sessions. Error bars indicate SEM. **d)** The performance improvement due to sleep (Performance) for Seq1(orange) and Seq2 (purple) in three cases: 1) Seq2 was trained as the same duration as Seq1 (*left*); 2) Seq2 was trained shorter than Seq1 (*middle*); (3) Seq2 was trained longer than Seq1 (right). In each case, we have three experiments: 1) Uncued, none of the sequences were cued during sleep; 2) Cued Seq1, only the cue associated with Seq1 was presented during sleep; 3) Cued Seq2, only the cue associated with Seq2 was presented during sleep. * p<0.05, ** p<0.01, *** p<0.001. N.S. represents no significant difference.

While both sequences revealed similar performance when they were trained for the same duration without presenting any cues during sleep, results changed when the sensory cue was applied during sleep (Fig. 8d, *left*). Here, we defined Δ*P* as the performance after sleep minus the performance before sleep, indicating the performance improvement due to sleep. If only the cue associated with Seq1 was presented during sleep, Δ*P* for Seq1 was significantly increased (Uncued vs. Cued Seq1: 17.3%±2.48% vs. 28.7%±3.20%, t(38)=2.877, p=0.0076, two-sample t test, Fig. 8d, *left*, orange bar), while for Seq2 was reduced but not significantly (Uncued vs. Cued Seq1: 17.2%±2.90% vs. 10.7%±3.38%, t(38)=-1.4614, p=0.1521, two-sample t test, Fig. 8d, *left*, purple bar). If only the cue associated with Seq2 was presented during sleep, for Seq2 was significantly increased (Uncued vs. Cued Seq2: 17.2%±2.90% vs. 27.3%±1.74%, t(38)=-2.9881, p=0.0049, two-sample t test, Fig. 8d, *left*, purple bar), while Δ*P* for Seq1 was reduced but not significantly (Uncued vs. Cued Seq2: 17.3%±2.48% vs. p0.3%±3.53%, t(38)=1.624, p=0.1127, two-sample t test, Fig. 8d, *left*, orange bar). These results suggest that presenting the cue associated with one specific memory sequence accelerates the consolidation of that sequence without significant interfering with the consolidation of another memory sequence.

We next used different training times for two sequences: Seq2 was trained less than Seq1 (Fig. 8d, *middle*). This difference remained after sleep in uncued condition (Fig. 8d, *middle*, Uncued). In such case, when the cue associated with Seq1 was presented during sleep, Δ*P* for Seq2 was significantly reduced compared with the uncued condition (Cued Seq1 vs. Uncued: −10.5%±1.97% vs. 4.1%±2.66%, t(38)=-4.4112, p=8.19* 10^-5^, two-sample t test, Fig. 8d, *middle*, purple bar) and even became negative, indicating that weaker memory of Seq2 was partially erased due to the strong reactivation of Seq1. When the cue associated with weaker Seq2 was presented, Δ*P* for Seq2 was significantly increased (Uncued vs. Cued Seq2: 4.1%±2.66% vs. 11.9%±2.21%, t(38)=-2.2537, p=0.03, two-sample t test, Fig. 8d, *middle*, purple bar), without significant impact on Seq 1 performance.

Lastly, we trained Seq2 longer than Seq1, therefore, the initial (before sleep) memory strength of Seq2 was higher than that of Seq1 (Fig. 8d, *right*). After sleep, in uncued condition, Seq 2 showed a very significant performance increase while Seq 1 improvement was relatively reduced (compare Fig. 8d, *right*, Uncued to Fig. 8d, *left*, Uncued) likely because of the interference between two sequences during sleep replay (Wei *et al.*, 2018). If only the cue associated with Seq1 was presented, Δ*P* for Seq1 was increased significantly but Δ*P* for Seq 2 was similar to that after uncued sleep. If only the cue associated with Seq2 was presented, Δ*P* for both Seq1 and Seq2 was also not significantly different compared with uncued sleep (Uncued vs. Cued Seq2: 30.7%±2.33% vs. 34%±2.58%, t(38)=-0.9483, p=0.349, two-sample t test, Fig. 8d, *right*, purple bar), indicating that consolidation of the stronger Seq2 does not have a room to further increase in a presence of a cue. Overall, these results suggest that (a) cues during slow-wave sleep are particularly effective to improve consolidation of the weak memories and (b) applying cues associated with one specific memory may lead to performance degradation for other, particularly weak memories, that are not cued.

## Discussion

In this study, using a realistic computational model of the thalamocortical network implementing sleep stages (Krishnan *et al.*, 2016) and synaptic plasticity (Wei, Krishnan and Bazhenov, 2016; Wei *et al.*, 2018), we aimed at developing a better understanding of how external stimulation during sleep can enhance memory consolidation. Our study suggests that both training-associated cues or auditory stimulation during SWS can promote spike sequence replay and enhance memory performance after sleep. The cues only strengthened the memory traces when delivered at or near the optimal phase - just before Down to Up state transition. This same phase relationship also applied for auditory stimulation during sleep, not previously related to the learning context. In both cases consolidation was improved because stimulation was able to shape the spatio-temporal pattern of sleep slow waves to promote Up state initiation near the site when trained memory was encoded. When multiple memories were trained and one memory was associated with a specific cue during training, sensory cues during SWS could selectively strengthen the associated memories but could also lead to performance degradation for other, particularly weak, memories.

### The mechanisms of strengthening memories by stimulation during NREM sleep

Synaptic plasticity is believed to be the cellular mechanism of learning and memory in the brain (Ho, Lee and Martin, 2011). Evidence is accumulating that recent memories are consolidated during NREM sleep (Rasch and Born, 2013; Born and Wilhelm, 2012; Diekelmann and Born, 2010a; Walker and Stickgold, 2004) through replay of sequences of cell-firing patterns that occur during waking (Ji and Wilson, 2007a; Euston, Tatsuno and McNaughton, 2007; Peyrache *et al.*, 2009b). This replay is thought to be orchestrated by hippocampal and thalamocortical patterns of activity (Maingret *et al.*, 2016; Girardeau *et al.*, 2009; Ego-Stengel and Wilson, 2010; Marshall *et al.*, 2006; Ngo *et al.*, 2013). We recently showed, using computer models, that the sequences of the cortical neurons’ firing trained in awake, are replayed spontaneously during NREM sleep; this enhances the synaptic connections associated with the trained memory resulting in memory improvement (Wei *et al.*, 2018). In this present study, we explored how external stimulation augments an internal and naturally occurring mechanism of memory reactivation, resulting in an enhanced memory consolidation process.

Recently, empirical studies revealed a powerful new tool to augment memory consolidation process - targeted memory reactivation (TMR) during sleep (for review, see (Oudiette and Paller, 2013; Schouten *et al.*, 2017)). In these experiments, stimulation (such as sounds or odors) presumably enhance reactivation of the relevant neuronal representations that improves memory consolidation. TMR was shown to improve both hippocampus-independent procedural memories (Antony *et al.*, 2012; Schonauer, Geisler and Gais, 2014; Cousins *et al.*, 2014; Cousins *et al.*, 2016) and hippocampus-dependent declarative memories (Rudoy *et al.*, 2009; Rasch *et al.*, 2007; Batterink, Creery and Paller, 2016; Oyarzun *et al.*, 2017). In one experiment, an odor was associated with learning a spatial card location task; subsequent presentation of the odor cue during sleep led to selective enhancement of this memory compared to the sleep without cueing (Rasch *et al.*, 2007). Similarly, auditory cues that were paired with visual input during learning, enhanced memory consolidation when presented during subsequent sleep (Rudoy *et al.*, 2009). The timing of the stimulation with respect to ongoing sleep slow oscillation determined the effect of a cue on memory consolidation (Batterink, Creery and Paller, 2016; Santostasi *et al.*, 2016). Specifically, the closed-loop stimulation presented during Down state, just before transition from Down to Up state of slow oscillation, had high impact on memory consolidation compared to the other stimulation phase. For overlapping memories, whether the auditory cues strengthen or weaken a memory also depended on the memory strength (Oyarzun *et al.*, 2017). Notably, the beneficial effect of external reactivation only occurred during sleep. Presenting the same cues during wakefulness was not effective for either declarative memories (Rasch *et al.*, 2007) or procedural memories (Schonauer, Geisler and Gais, 2014).

We recently reported, using biophysical models of the thalamocortical system, that sleep reactivation occurs during Up states of the slow oscillation (Wei *et al.*, 2018). When synaptic connectivity matrix was modified in awake state by sequence training, the trained sequences were spontaneously replayed during Down to UP state transitions and also during early phase of the Up states leading to the further changes of synaptic weights (consolidation). For memories represented by the sequences of neuronal activation (as in this study), beginning of a sequence was commonly also a site where local spontaneous Up state was initiated. In this new study, we showed that the effect of stimulation on memory replay depends on the ability of stimuli to affect, at least locally, the spatio-temporal pattern of sleep slow waves. When a local stimulus was delivered at the “correct” phase of the slow oscillation (during Down state but close to transition to Up state) and to the “correct” network location (near the trained sequence), it effectively triggered local Up state initiation and promoted replay of a sequence associated with this stimulation site. Stimuli delivered at “wrong” timing, e.g., when the network is already active or just transited to the Down state, or “wrong” location, e.g., far away from the beginning of a sequence in question, could not facilitate replay of that sequence.

### Closed-loop auditory stimulation during sleep: local or broad cue?

Different types of the auditory stimulation, represented either by the learning-specific cues (Batterink, Creery and Paller, 2016) or pink noise (Ngo *et al.*, 2015; Ngo *et al.*, 2013; Santostasi *et al.*, 2016), are all able to enhance memory consolidation. We found, however, that the optimal phase and the mechanisms were somewhat different for these two different types of sensory stimulation.

For the learning-specific cue, we hypothesized that the cue only affects a small population of neurons (“local cue”). This may explain why stimulation is only effective *in vivo* when it was applied at specific phase of the slow waves (Batterink, Creery and Paller, 2016). In our study, we identified that the optimal phase depends on the influence of the stimuli on the Up state initiation. When stimulus is delivered just before Down to Up state transition when the network is sensitive to external perturbations, the stimulus affects (possibly locally) the spatiotemporal pattern of slow waves promoting the sequence replay and increase of synaptic weights associated with trained sequence in question.

For the pink noise auditory stimulation, stimulation can potentially activate a broad group of neurons (“broad cue”). It was shown in vivo that such broad stimuli could enhance memory consolidation by enhancing the slow oscillation (Ngo *et al.*, 2013; Ngo *et al.*, 2015). Importantly, the pink auditory stimulation was effective *in vivo* when the sound occurs in synchrony with the slow oscillation Up states (Ngo *et al.*, 2013). In the model, broader stimulation led to higher increase in memory performance compare to the local one. Also, while the peak performance was still achieved for stimuli delivered at the network transition from Down to Up state, the difference was smaller between optimal and suboptimal stimulations. In all cases increase in performance was linked to a significant increase in amplitude and synchrony of slow oscillation, in agreement with in vivo data. Considering that in vivo the slow oscillation is not necessarily well synchronized across entire cortex (and definitely less synchronized than that in our model) (Nir *et al.*, 2011), we predict that for broader stimuli the optimal stimulation would be during the early phase of an Up state when it is most capable to induce high amplitude synchronized slow wave across large cell populations of the cortical network.

Together, our study provides insight into how the memory consolidation may be affected by stimulation applied during slow-wave sleep. We consider stimulation both as a tool to manipulate the sleep rhythms to understand mechanisms behind the role of sleep in memory consolidation and as an approach to develop clinical interventions with a goal of enhancing memory and learning. The auditory stimulation during sleep has the potential application to enhance the sleep-dependent effects on memory in cognitively healthy human subjects and in those with amnestic mild cognitive impairment.

## ACKNOWLEDGEMENTS

This work was supported by National Science Foundation (IIS-1724405), DARPA (HR0011-18-2-0021), NIH (RF1MH117155), ONR (N00014-16-1-2829-P00005)

